# Targeted Genome Editing of Bacteria Within Microbial Communities

**DOI:** 10.1101/2020.07.17.209189

**Authors:** Benjamin E. Rubin, Spencer Diamond, Brady F. Cress, Alexander Crits-Christoph, Christine He, Michael Xu, Zeyi Zhou, Dylan C. Smock, Kimberly Tang, Trenton K. Owens, Netravathi Krishnappa, Rohan Sachdeva, Adam M. Deutschbauer, Jillian F. Banfield, Jennifer A. Doudna

## Abstract

Knowledge of microbial gene functions comes from manipulating the DNA of individual species in isolation from their natural communities. While this approach to microbial genetics has been foundational, its requirement for culturable microorganisms has left the majority of microbes and their interactions genetically unexplored. Here we describe a generalizable methodology for editing the genomes of specific organisms within a complex microbial community. First, we identified genetically tractable bacteria within a community using a new approach, Environmental Transformation Sequencing (ET-Seq), in which non-targeted transposon integrations were mapped and quantified following community delivery. ET-Seq was repeated with multiple delivery strategies for both a nine-member synthetic bacterial community and a ~200-member microbial bioremediation community. We achieved insertions in 10 species not previously isolated and identified natural competence for foreign DNA integration that depends on the presence of the community. Second, we developed and used DNA-editing All-in-one RNA-guided CRISPR-Cas Transposase (DART) systems for targeted DNA insertion into organisms identified as tractable by ET-Seq, enabling organism- and locus-specific genetic manipulation within the community context. These results demonstrate a strategy for targeted genome editing of specific organisms within microbial communities, establishing a new paradigm for microbial manipulation relevant to research and applications in human, environmental, and industrial microbiomes.

---

Genetic mutation and observation of phenotypic outcomes are the primary means of deciphering gene function in microorganisms. This classical genetic approach requires manipulation of isolated species, limiting knowledge in three fundamental ways. First, the vast majority of microorganisms have not been isolated in the laboratory and are thus largely untouched by molecular genetics^1^. Second, emergent properties of microbial communities that may arise due to interactions between their constituents, remain mostly unexplored^2^. Third, microorganisms grown and studied in isolation quickly adapt to their new lab environment, obscuring their true “wild type” physiology^3^. Since most microorganisms relevant to the environment, industry and health live in communities, approaches for precision genome modification (editing) in community contexts will be transformative.

Advances toward genome editing within microbial communities have included assessing gene transfer to microbiomes using selectable markers^4–9^, microbiome manipulation leveraging pre-modified isolates^10^, and use of temperate phage for species-specific integration of genetic payloads^11^. However, a generalizable strategy for programmable organism- and locus-specific editing within a community of wild-type microbes has not yet been reported^12^.

Here we show that specific organisms within microbial communities can be targeted for site-specific genome editing, enabling manipulation of species without requiring prior isolation or engineering. Using a new method developed for this study, Environmental Transformation Sequencing (ET-Seq), we identified genetically accessible species within a nine-member synthetic community and among previously non-isolated species in a 197-member bioremediation community. These results enabled targeted genome editing of microbes in the nine-member community using DNA-editing All-in-one RNA-guided CRISPR-Cas Transposase (DART) systems developed here. The resulting species-specific editing provides the first broadly applicable strategy for organism- and locus-specific genetic manipulation within a microbial community, hinting at new emergent properties of member organisms and methods for controlling microorganisms within their native environments.

## ET-Seq detects genetically accessible microbial community members

Editing organisms within a complex microbiome requires knowing which constituents are accessible to nucleic acid delivery and editing. We developed ET-Seq to assess the ability of individual species within a microbial community to acquire and integrate exogenous DNA (Fig. 1a). In ET-Seq, a microbial community is exposed to a randomly integrating mobile genetic element (here, a *mariner* transposon), and in the absence of any selection, total community DNA is then extracted and sequenced using two protocols. In the first, we enrich and sequence the junctions between the inserted and host DNA to determine insertion location and quantity in each host. This step requires comparison of the junctions to previously sequenced community reference genomes. In the second, we conduct low-depth metagenomic sequencing to quantify the abundance of each community member in a sample (Extended Data Fig. 1a). Together these sequencing procedures provide relative insertion efficiencies for microbiome members. To normalize these data according to a known internal standard, we add to every sample a uniform amount of genomic DNA from an organism that has previously been transformed with, and selected for, a *mariner* transposon. In the internal standard, we expect every genome to contain an insertion. The final output of ET-Seq is a fractional number representing the proportion of a target organism’s population that harbored transposon insertions at the time DNA was extracted, or insertion efficiency (Extended Data Fig. 1b). To facilitate the analysis of these disparate data, we developed a complete bioinformatic pipeline for quantifying insertions and normalizing results according to both the internal control and metagenomic abundance (https://github.com/SDmetagenomics/ETsuite and Methods). Together the experimental and bioinformatic approaches of ET-Seq reveal species-specific genetic accessibility by measuring the percentage of each member of a given microbiome that acquires a transposon insertion.

**Fig. 1.**
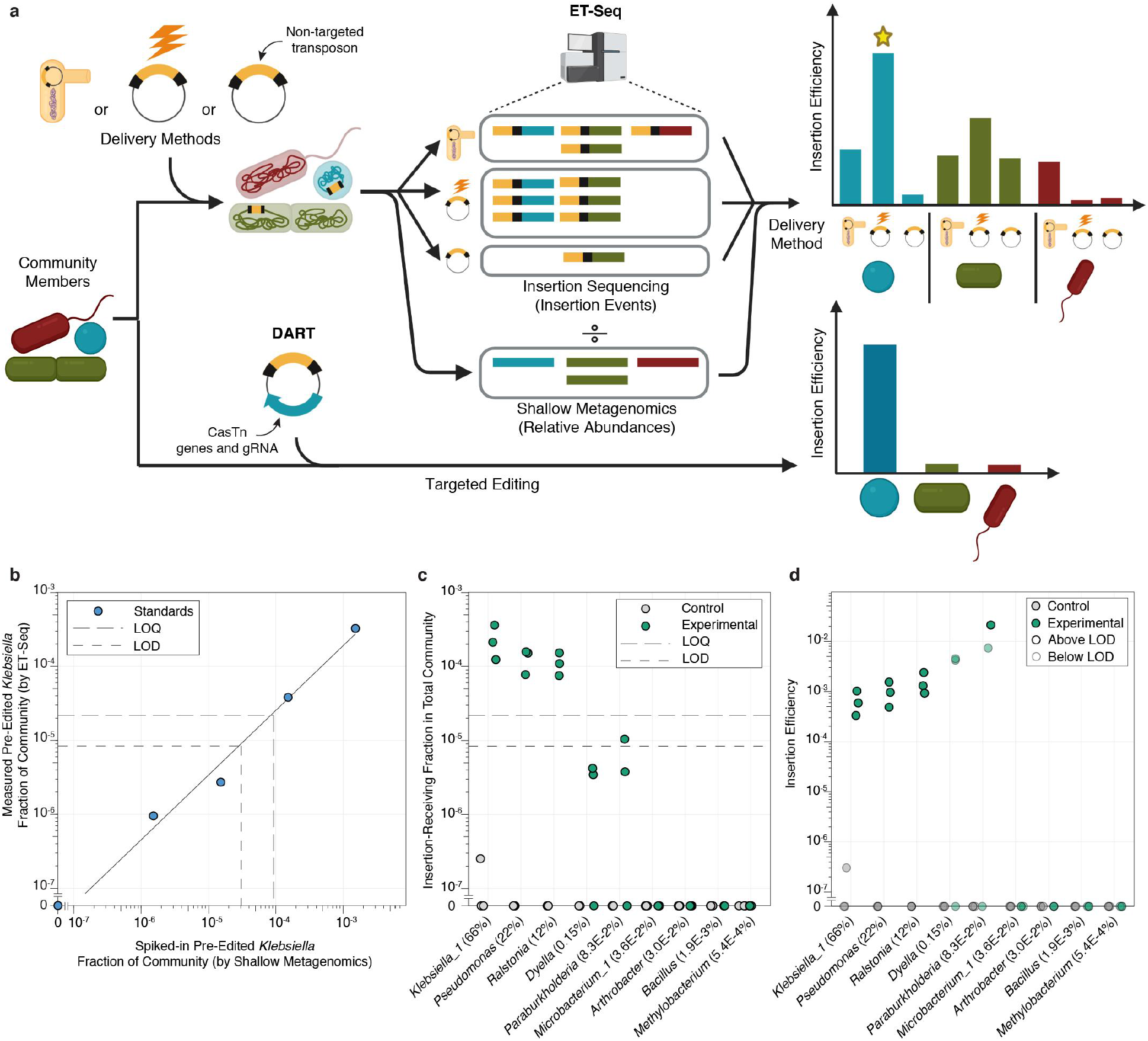
ET-Seq for quantitative measurement of insertion efficiency in a microbial community. **a**, ET-Seq provides data on insertion efficiency of multiple delivery approaches, including conjugation, electroporation, and natural DNA transformation, on microbial community members. In this illustrative example, the blue strain is most amenable to electroporation (star). This data allows for the determination of feasible targets and delivery methods for DART targeted editing. **b,** ET-Seq determined efficiencies for known quantities of spiked-in pre-edited *K. michiganensis* (*Klebsiella_1*). Data shown is the mean of three technical replicates. LOD is the lowest insertion fraction at which accurate detection of insertions is expected and LOQ is the lower limit at which this fraction is expected to be quantifiable. Solid line is the fit of the linear regression to the data not including zero (n = 4 independent samples) that is used to calculate LOD and LOQ (slope = 0.2137; intercept = 1.813*10^-6^). **c-d,** ET-Seq determined insertion efficiencies in the nine-member consortium (n = 3 biological replicates) with conjugative delivery shown as **c,** a portion of the entire community and **d,** a portion of each species. Control samples received no exogenous DNA. Average relative abundances of community constituents across conjugation samples (n = 6 independent samples) are indicated in parentheses. LOD and LOQ are indicated in plots by short and long dashed lines respectively.

ET-Seq was developed and tested on a nine-member microbial consortium made up of bacteria from three phyla that are often detected and play important metabolic roles within soil microbial communities (Supplementary Table 1). We initially endeavored to test the accuracy and detection limit by adding to the nine-member community a known amount of a previously prepared *mariner* transposon library of one of its member species, *Klebsiella michiganensis* M5a1 (*Klebsiella_1*). The ET-Seq derived insertion efficiencies were closely correlated to the known fractions of edited *K. michiganensis* present in each sample (Fig. 1b). Using this data we calculated a limit of detection (LOD) and limit of quantification (LOQ) for our approach (Methods). The LOD suggests that a fraction of ≥ 8.4*10^-6^ of transformed cells out of the total community would be detectable by ET-Seq.

Next, the *mariner* transposon vector was delivered to the nine-member community through conjugation. We could measure conjugation reproducibly and quantitatively in the three species that grew to make up over 99% of the community (Fig. 1c). We further normalized insertion efficiency in each species according to its abundance so that their insertion efficiencies represent insertion containing cells as a portion of total cells for each species (Fig. 1d). Even for *Paraburkholderia caledonica*, which made up ~0.1% of the community, we could measure insertions. We detected no insertions in the remaining community members, which was expected given their extreme rarity in the community (less than 0.05%).

We next used ET-Seq to compare insertion efficiencies in the nine-member community after transposon delivery by conjugation, natural transformation with no induction of competence, or electroporation of the transposon vector. Together these approaches showed reproducible insertion efficiencies above the LOD in five of the nine community members (Fig. 2a and Extended Data Fig. 2). Additionally we could identify preferred delivery methods for some members in this community context, such as electroporation being consistently reproducible for *Dyella japonica UNC79MFTsu3.2* while conjugation was not. These results show that ET-Seq can identify and quantify genetic manipulation of microbial community members and reveal suitable DNA delivery methods for each.

**Fig. 2.**
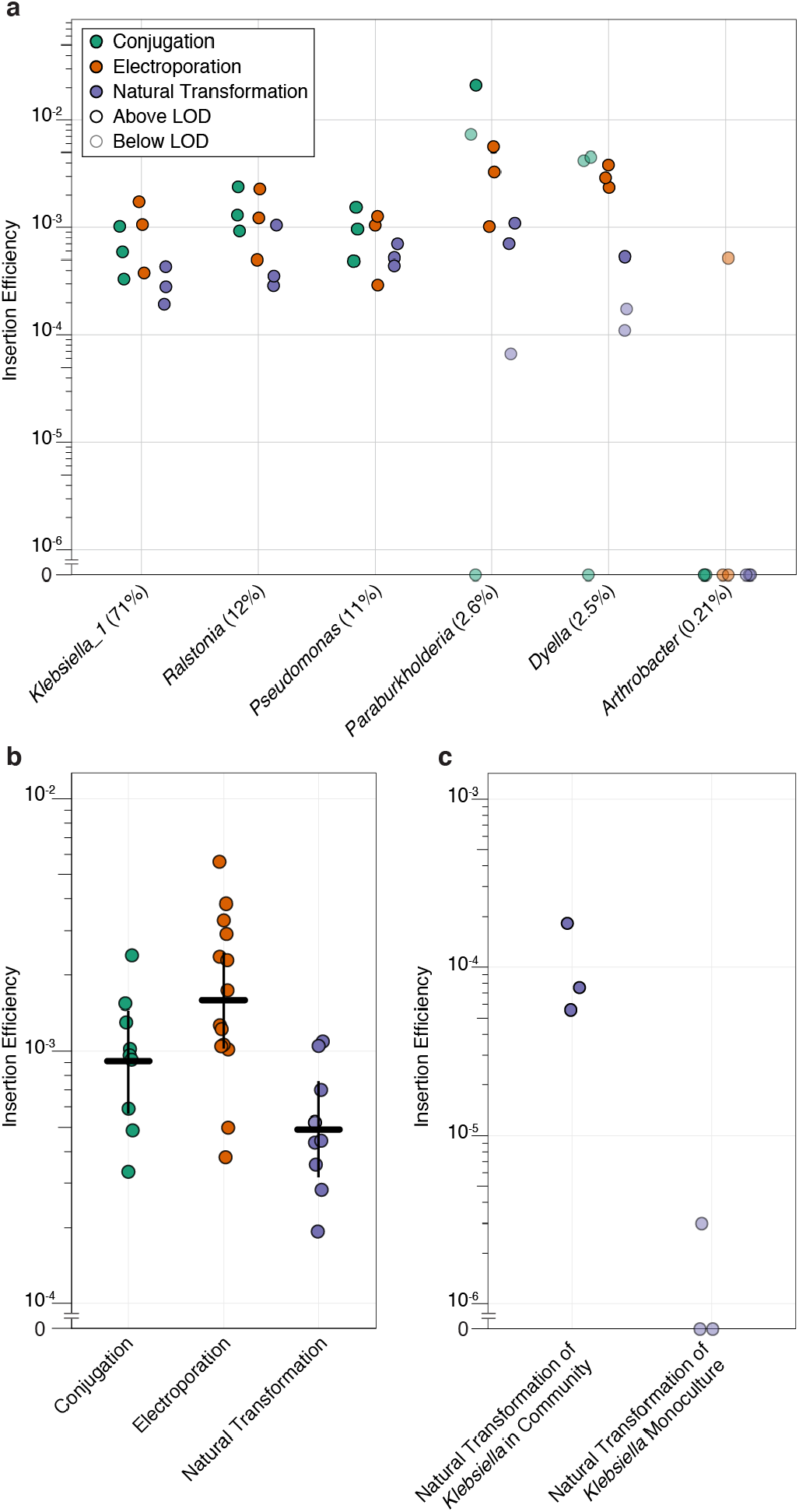
ET-Seq detection of insertion efficiency across multiple delivery approaches. **a**, ET-Seq determined insertion efficiencies for conjugation, electroporation, and natural transformation on the nine-member consortium (n = 3 biological replicates). Only members with at least one positive insertion efficiency value across the delivery methods are shown. Average relative abundance across all samples (n = 18 independent samples) is indicated in parentheses. **b,** Comparing delivery strategies across data from all organisms. Cross bars indicate the mean value and whiskers denote the 95% confidence interval for the mean (Conjugation n = 9; Electroporation n = 14; Natural Transformation n = 9). **c**, Comparison of insertion efficiencies measured for natural transformation of *K. michiganensis* (*Klebsiella_1*) in isolated culture (n = 3 biological replicates) compared to *K. michiganensis* grown in the community context (n = 3 biological replicates).

Notably, five organisms exhibited some degree of natural competency, although average efficiencies were significantly lower for natural transformation than for delivery through electroporation (ANOVA followed by Tukey’s HSD; two-sided test; p = 0.0009) (Fig. 2b). To the best of our knowledge, no isolates of the *Klebsiella* genus including *K. michiganensis* are known to be naturally competent. We conducted a second experiment to compare the insertion efficiency of *K. michiganensis* cultivated in isolation versus grown in the community context. ET-Seq returned no values above the LOD for natural transformation of *K. michiganensis* in isolation, but within the community ET-Seq returned values well into the quantifiable range (Fig. 2c). This apparent emergent property of natural competence within a small synthetic community provides tantalizing support for the possibility of community induced natural transformation, an idea suggested in previous work, but experimentally unstudied due to lack of tools for measuring horizontal gene transfer events within a community^13,14^.

## Genetic accessibility of uncultivated species within an environmental microbiome

To test the potential for editing in a complex and environmentally realistic community that has not been reduced to isolates, we conducted ET-Seq on a genomically characterized 197 member bioreactor-derived consortium that degrades thiocyanate (SCN^-^), a toxic byproduct of gold processing^15^. We sampled biofilm from the reactor and conducted ET-Seq with a panel of delivery techniques: conjugation, electroporation, and natural transformation. Across ET-Seq replicates at least one measurement above the LOD was identified for 15 members of the bioreactor community. We also note that the transformed organisms make up ~87% of the bacterial fraction by relative abundance (Fig. 3a, Extended Data Fig. 3). Ten of these organisms were species that had not previously been isolated or edited (Supplementary Table 1), and overall members from 7 of the 12 phyla detected in this consortium were successfully transformed (Fig. 3b). This included an *Afipia sp*. known to play an important role in the thiocyanate degradation process. Additionally, one of the transposon recipients, *Microbacterium ginsengisoli* (*Microbacterium_3*), is a putative host for Saccharibacteria, a candidate phyla radiation (CPR) organism that has been observed in this reactor system^16^. Notably, members of the CPR are resistant to typical isolation techniques due to heavy dependence on other community members, and little is known about the nature of their likely symbiotic relationships with other organisms^17^. Here, ET-Seq has uncovered a genetically tractable putative host organism, raising the possibility of genetically editing the host to probe CPR/host symbiotic relationships within a complex microbial community. In this way, ET-Seq reveals genetic accessibility and the tools necessary to achieve it in previously unapproachable and biologically important members of an environmentally relevant community.

**Fig. 3.**
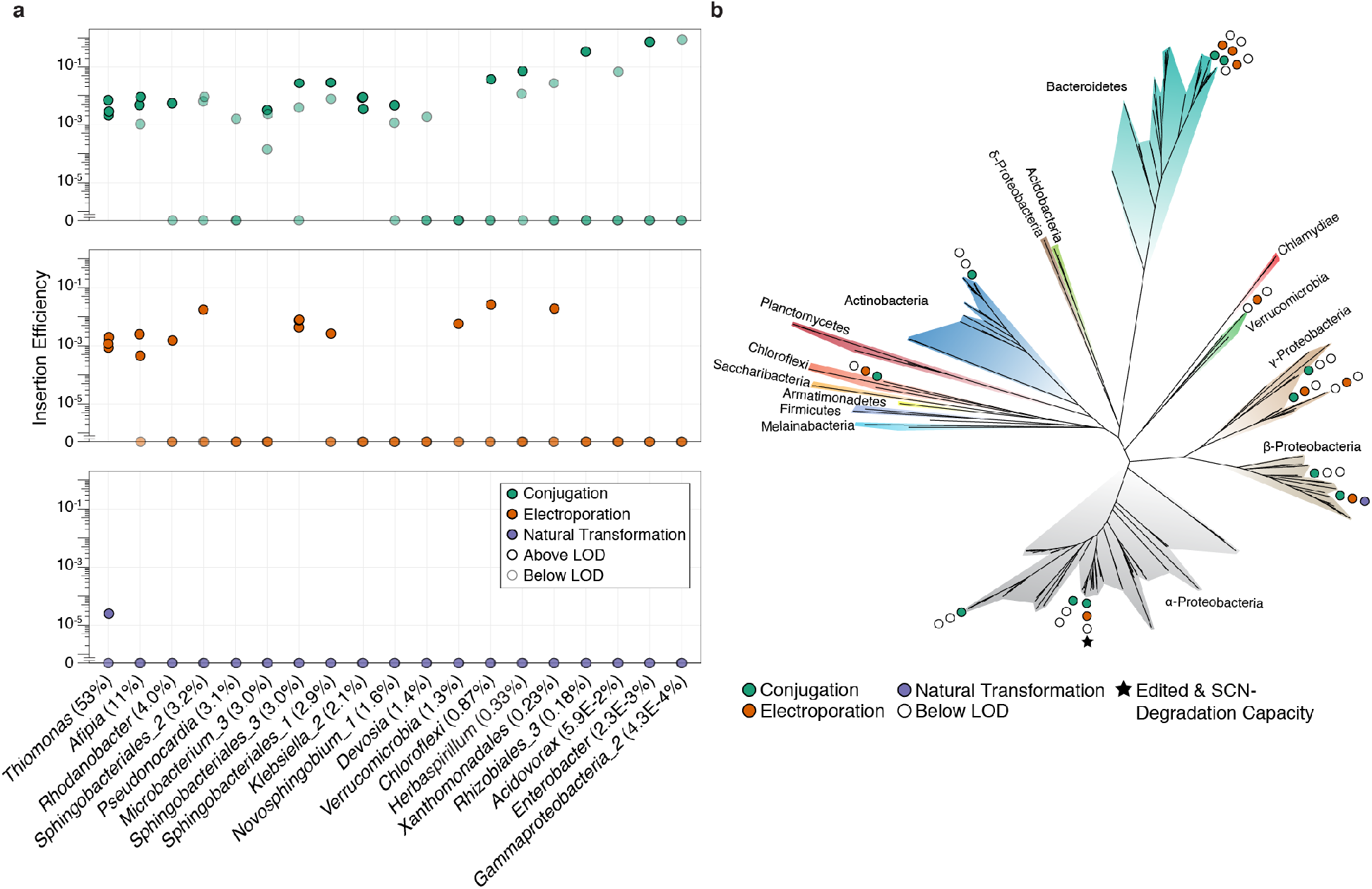
ET-Seq detection of insertion efficiency in thiocyanate-degrading bioreactor. **a,** ET-Seq determined insertion efficiencies for conjugation, electroporation, and natural transformation on the thiocyanate-degrading bioreactor community (n = 3 biological replicates). Average relative abundance across all samples is indicated in parentheses (n = 17 independent samples). **b,** A ribosomal protein S3 (rpS3) phylogenetic tree of all organisms in the thiocyanate-degrading bioreactor genome database. Only community members receiving at least one insertion by conjugation, electroporation, or natural transformation detectable above LOD are indicated by filled circles. Filled circles indicate success of method and open circles indicate method was not detected. A star indicates genomically encoded SCN^-^ degradation capacity in organisms shown to be edited by at least one method. Tree was constructed from an alignment of 262 rps3 protein sequences using IQ-TREE.

## Targeted genome editing in microbial communities using CRISPR-Cas transposases

The ability to introduce genome edits to a single type of organism in a microbial community and to target those edits to a defined location within its genome would be a foundational advance in microbiological research and would have many useful applications. We reasoned that RNA-guided CRISPR-Cas Tn7 transposases could provide the ability to both ablate function of targeted genes and deliver customized genetic cargo in organisms shown to be genetically tractable by ET-Seq^18–20^ (Fig. 1a). However, the two-plasmid ShCasTn^18^ and three-plasmid VcCasTn^19^ systems are not amenable to efficient delivery within complex microbial communities or even beyond *E. coli* due to their multiple plasmids. Since ET-Seq identified conjugation and electroporation as broadly effective delivery approaches in the tested communities, we designed and constructed all-in-one conjugative versions of these CasTn vectors that could be used for delivery by either strategy (Fig. 4a and Methods). These DART systems are barcoded and compatible with the same sequencing methods used for ET-Seq, and can be used to assay the efficacy of CRISPR-Cas-guided transposition into the genome of a target organism.

**Fig. 4.**
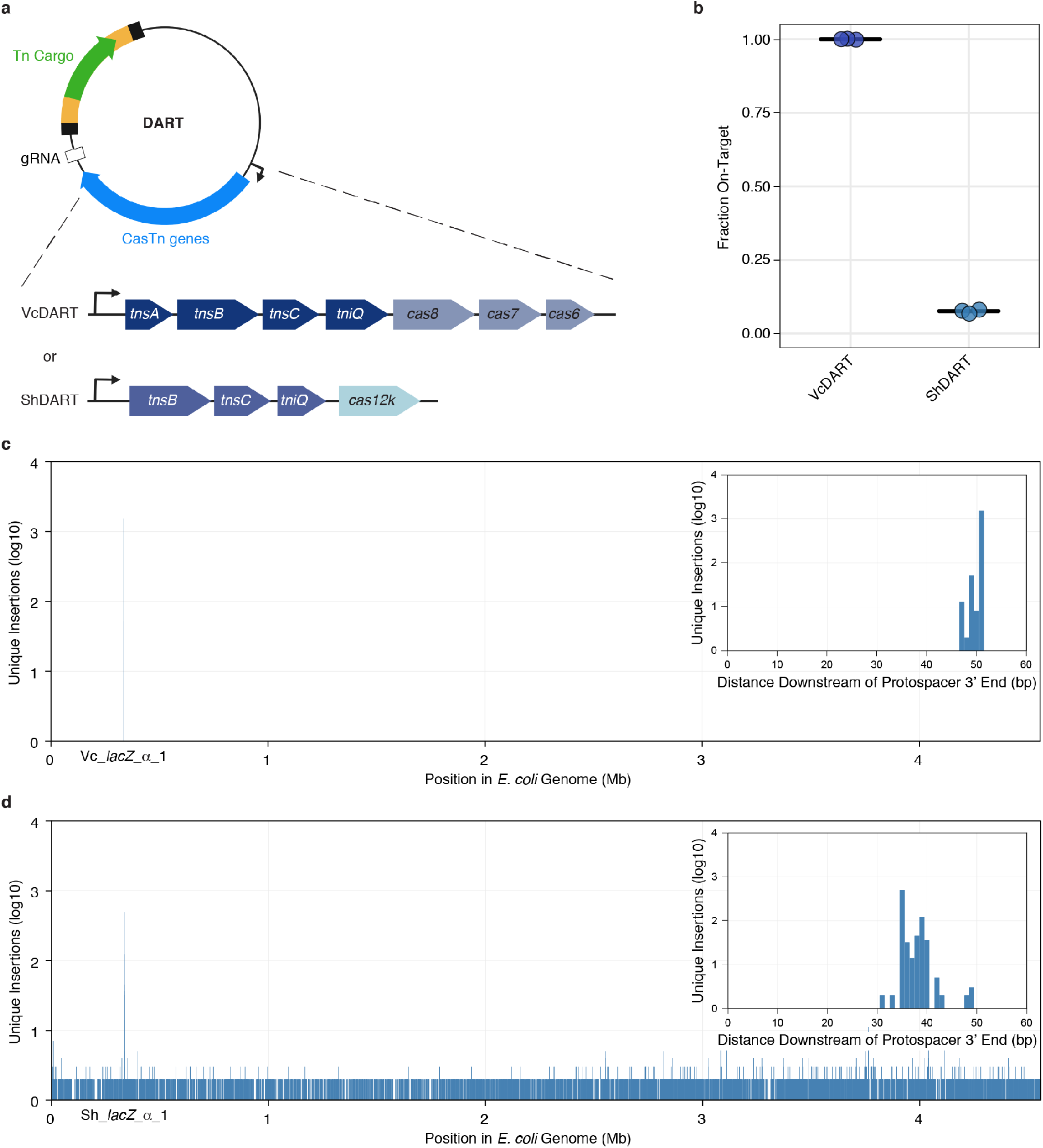
Benchmarking all-in-one conjugal targeted vectors. **a,** Schematic of VcDART and ShDART delivery vectors. **b,** Fraction of insertions that occur in a 60 bp window around the target site. Mean for three independent biological replicates is shown as cross bars. **c-d,** Aggregate unique insertion counts (n = 3 biological replicates) across the *E. coli* BL21(DE3) genome, determined by presence of unique barcodes, using **c,** VcDART and **d,** ShDART. The inset shows a 60 bp window downstream of the target site. Insertion distance downstream of the target site is calculated from the 3’ end of the protospacer.

We compared the transposition efficiency and specificity of the DART systems in *E. coli* in order to select the most promising candidate for targeted genome editing in microbial communities. VcDART and ShDART systems harboring Gm^R^ cargo with a *lacZ*-targeting or non-targeting guide RNA were conjugated into *E. coli* to quantify transposition efficiency, and target site specificity was assayed using ET-Seq following outgrowth of transconjugants in selective medium (Methods and Extended Data Fig. 4a). While ShDART yielded approximately tenfold more colonies possessing insertions than ShDART (Extended Data Fig. 4), >92% of the selectable colonies obtained using ShDART were off-target, compared to no detectable off-target insertions for VcDART (Fig. 4b-d). Due to VcDART’s high target site specificity and the undesirable propensity for ShCasTn to co-integrate its donor plasmid^21,22^, we focused on VcDART to test the potential for targeted microbial community genome editing.

## Targeted microbial community editing by programmable transposition

We reasoned that RNA-programmed transposition could be deployed for targeted editing of specific types of organisms within a microbial consortium. ET-Seq had shown two species within the nine-member community, *K. michiganesis* and *Pseudomonas simiae* WCS417, to be both abundant and tractable by conjugation (Fig. 1d). We targeted both of these organisms using conjugation to introduce the VcDART vector into the community with guide RNAs specific to their genomes (Fig. 5a). Insertions were designed to produce loss-of-function mutations in the *K. michiganesis* and *P. simiae pyrF* gene, an endogenous counterselectable marker allowing growth in the presence of 5-fluoroorotic acid (5-FOA) when disrupted. The transposons carried two antibiotic resistance markers conferring resistance to streptomycin and spectinomycin (*aadA*) and carbenicillin (*bla*). Together the simultaneous loss-of-function and gain-of-function mutations allowed for a strong selective regime. VcDART targeted to *K. michiganensis* or to *P. simiae pyrF* followed by selection led to enrichment of these organisms to ~98% and ~97% pure culture respectively (Fig. 5b). No outgrowth was detected when using a guide RNA that did not target these respective microbial genomes. Recovered transformant colonies of *K. michiganensis* and *P. simiae* analyzed by PCR and Sanger sequencing showed full length, *pyrF*-disrupting VcDART transposon insertions 48-49 bp downstream of the guide RNA target site, consistent with CRISPR-Cas transposase-catalyzed transposition events at the desired genomic location (Fig. 4c and 5c). These results demonstrate that targeted genome editing using DART enables genetic manipulation of distinct members of a complex microbial community. This targeted editing of microorganisms in a community context can also enable subsequent exploitation of modified phenotypes.

**Fig. 5.**
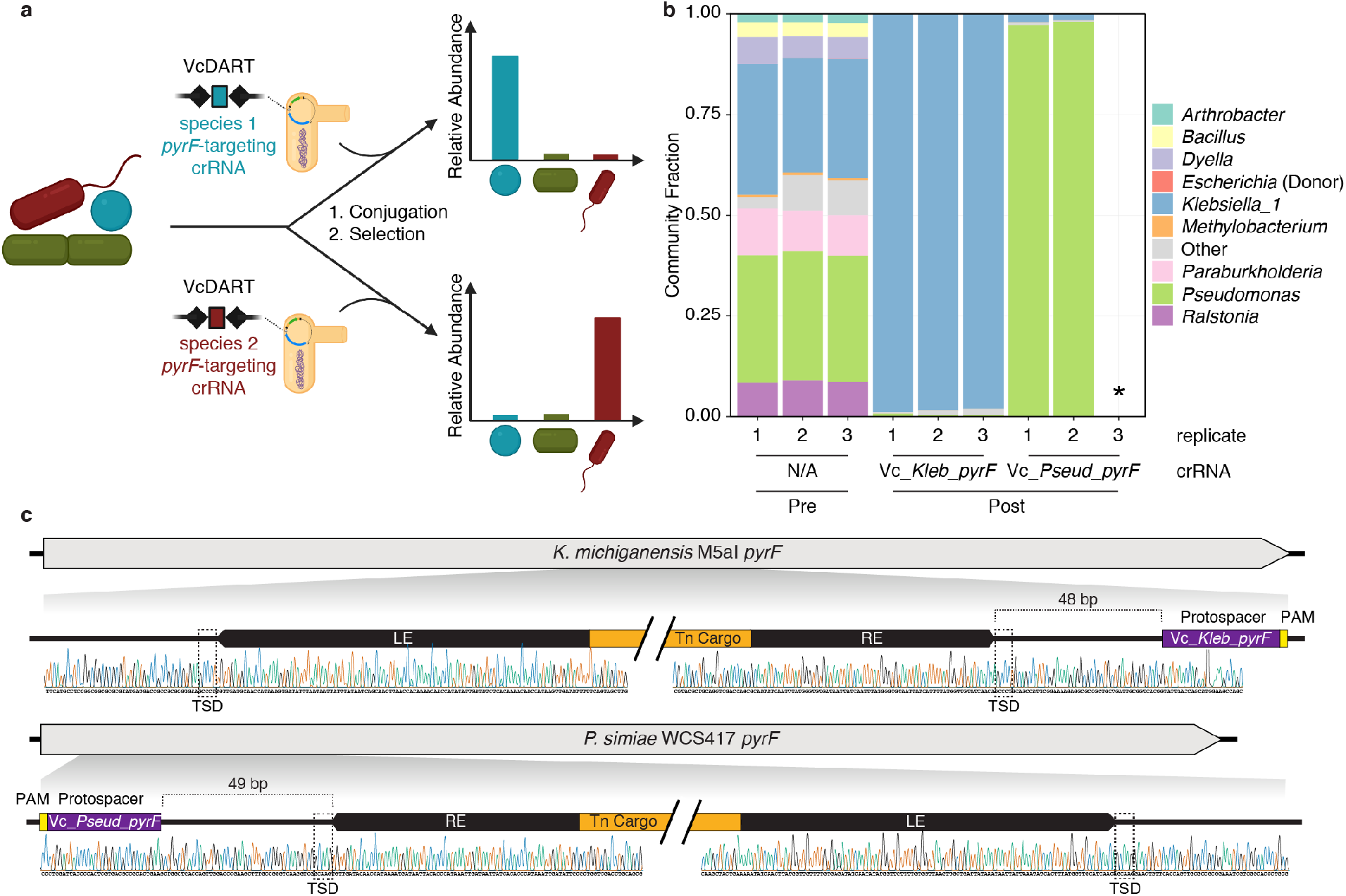
Targeted editing in the nine-member consortium. **a,** Conjugative VcDART delivery into a microbial community using species-specific crRNA, followed by selection for transposon cargo, facilitates selective enrichment of targeted organisms. **b,** Relative abundance of nine-member community constituents measured by 16S rRNA sequencing before conjugative VcDART delivery and after selection for *pyrF*-targeted transposition in *K. michiganensis* or *P. simiae* (n = 3 biological replicates). * indicates no growth detected in selective medium. **c,** Representative Sanger sequencing chromatogram of PCR product spanning transposon insertion site at targeted *pyrF* locus in *K. michiganensis* (top) and *P. simiae* (bottom) colonies following VcDART-mediated transposon integration and selection. Target-site duplications (TSD) are indicated with dashed boxes.

## Discussion

We have demonstrated organism- and locus-specific genome editing within a microbial community, providing a new approach to microbial genetics and microbiome manipulation for research and applications. ET-Seq revealed the genetic accessibility of organisms growing within microbial communities, including ten microbial species that had not been previously isolated or found to be genetically manipulated. The creation of all-in-one vectors encoding two naturally occurring CRISPR-Cas transposon systems enabled comparison of their targeted genome editing capabilities. These experiments showed that only one of the two systems, which we termed VcDART, enabled precise RNA-programmable microbial genome editing. The ability to conduct targeted genome editing of two bacteria within a nine-member synthetic community and to use the introduced genetic changes as a means of organism isolation demonstrates a new approach to microbiome manipulation. Traditionally, the combined steps of culturing an environmental microbe, determining the ideal means to transform it, and implementing targeted editing could take years or could fail altogether^23^. ET-Seq combined with VcDART compresses the pipeline for establishing genetics in microorganisms to weeks and expands the diversity of organisms that can be targeted beyond those that can be cultivated in isolation.

In addition to providing a roadmap for targeted microbial genome editing, ET-Seq can be used to discover and analyze horizontal gene transfer in complex communities. In this study, we observed unexpected community-dependent natural transformation in the nine-member community and characterized the horizontal gene transfer events experimentally in the complex microbiome of a thiocyanate-degrading bioreactor. In future experiments, multiple ET-Seq time points could be taken in a community after delivery to measure directly both the portion of the community receiving DNA and the persistence of gene transfer, rather than tracking horizontal gene transfer using bioinformatics^24^ or indirect experimental methods^4–9^. Furthermore, as ET-Seq is applied in increasingly diverse and complex environments, an atlas of editable taxa can be created including optimal delivery approaches. To expand this dataset, we plan to apply ET-Seq to new microbial communities being sampled for metagenomic sequencing, a natural pairing because ET-Seq depends on availability of genome sequences for the component organisms.

In the future, tools are needed to create more generally applicable and persistent targeted genome edits. The canonical approach involving antibiotic selection for edited bacteria is infeasible in large complex communities, where many natural resistances exist^25,26^. Even in the nine-member synthetic community used in this study, three antibiotics and counterselection were necessary to achieve strict selection. Improved delivery strategies and alternative positive and counter selection methods should enable more efficient editing. In the gut microbiome, porphyran has been used successfully for selection of a spiked-in organism capable of utilizing this compound^27^. Such approaches for more efficient microbial community editing will enable research to answer fundamental questions as well as allow manipulation of agricultural, industrial, and health-relevant microbiomes. The combination of ET-Seq and DART systems presented here provide the foundation of the new field of *in situ* microbial genetics.

## Methods

### Plasmid construction and barcoding

For ET-Seq measurement of genetic tractability in community members, DNA encoding a non-targeted *mariner* transposon was delivered. The *mariner* transposon integrates into “TA” sequences in recipient cells. For delivery of the *mariner* transposon, we made use of the previously created pHLL250 vector, which contains an RP4 origin of transfer (oriT), AmpR, conditional (*pir*^+^-dependent) R6K origin, and an AseI restriction site to facilitate depletion of vector from DNA samples in ET-Seq library preparations^1^. Unique to each transposon on this vector is a random 20 bp barcode sequence to aid in the discrimination of unique insertion events from duplications of the same insertion due to cell division or PCR.

DART vectors were designed to encode all components required for delivery and editing (Supplementary Table 2 and Extended Data Fig. 4). VcCasTn genes, crRNA, and Tn were synthesized as gBlocks (IDT). pHelper_ShCAST_sgRNA was a gift from Feng Zhang (Addgene plasmid #127921; http://n2t.net/addgene:127921; RRID:Addgene_127921) and was used to clone ShCasTn genes and sgRNA. pDonor_ShCAST_kanR was a gift from Feng Zhang (Addgene plasmid # 127924; http://n2t.net/addgene:127924; RRID:Addgene_127924) and was used to clone the ShCasTn transposon. *tns* genes, *cas* genes, and crRNA/sgRNA were consolidated into a single operon (with various promoters and transcriptional configurations) on the same vector as the cognate transposon. The left end of the cognate Tn was encoded downstream of the crRNA/sgRNA, followed by Tn cargo, barcode, and Tn right end. DART Tn LE and RE were designed to include the minimal sequence that both included all putative TnsB binding sites and was previously shown to be functional^2,3^. Specifically, VcDART LE (108 bp) and RE (71 bp) each encompass three 20 bp putative TnsB binding sites, spanning from the edge of the 8 bp terminal ends to the edge of the third putative TnsB binding site^2^. ShDART LE (113 bp) spans the boundaries of the long terminal repeat and both additional putative TnsB binding sites, while the RE (211 bp) encompasses the long terminal repeat and all four additional putative TnsB binding sites^3^.

Vectors were cloned using BbsI (NEB) Golden Gate assembly of part plasmids, each encoding different regions of the final plasmid. Of note, the backbone encodes RP4 oriT, AmpR, conditional R6K origin, and an AsiSI+SbfI double digestion site for vector depletion during ET-Seq library preparations. A 2xBsaI spacer placeholder enabled spacer cloning with BsaI (NEB) Golden Gate. A 2xBsmBI barcode placeholder was encoded immediately inside the Tn right end and was used for barcoding as described below. Part plasmids were propagated in *E. coli* Mach1-T1R (QB3 Macro Lab). Golden Gate reactions for all-in-one vector assembly were purified with DNA Clean & Concentrator-5 (Zymo Research) and electroporated into *E. coli* EC100D-*pir*+ (Lucigen).

DART vectors were barcoded by BsmBI (NEB) Golden Gate insertion of random barcode PCR product into the 2xBsmBI barcode placeholder using a previously reported method^4^ with slight modifications. A 56-nt ssDNA oligonucleotide encoding a central tract of 20 degenerate nucleotides (oBFC1397) was amplified with BsmBI-encoding primers oBFC1398 and oBFC1399 using Q5 High-Fidelity 2X Master Mix (NEB) in a six-cycle PCR (98°C for 1 min; six cycles of 98°C for 10 s, 58°C for 30 s, and 72°C for 60 s; and 72°C for 5 min). Barcoding Golden Gate reactions were purified with DNA Clean & Concentrator-5. To remove residual non-barcoded vector, reactions were digested with 15 U BsmBI at 55°C for at least 4 hr, heat inactivated at 80°C for 20 min, treated with 10 U Plasmid-Safe ATP-Dependent DNase (Lucigen) exonuclease at 37°C for 1 hr, heat inactivated at 70°C for 30 min, and purified with DNA Clean & Concentrator-5.

Randomly barcoded conjugative vectors were electroporated into *E. coli* EC100D-*pir*+, followed 1 hr recovery in 1 mL pre-warmed SOC (NEB) at 37°C 250 rpm, serial dilution and spot plating on LB agar plus 100 μg mL^-1^ carbenicillin to estimate library diversity, and plating the full transformation across 5 LB agar plates containing carbenicillin (and other appropriate antibiotics when Tn cargo contained other resistance cassettes). To prepare barcoded conjugative vector plasmid stock, all 5 agar plates were scraped into a single pool and midiprepped (Zymo Research). All conjugations were performed using the diaminopimelic acid (DAP) auxotrophic RP4 conjugal donor *E. coli* strain WM3064. Donor strains were prepared by electroporation with 200 ng barcoded vectors, followed by recovery in SOC plus DAP at 37°C and 250 rpm and inoculation of the entire recovery culture into 15 mL LB containing DAP and carbenicillin in 50 mL conical tubes, followed by overnight cultivation at 37°C and 250 rpm. Donor serial dilutions were spot plated on LB agar plus carbenicillin to estimate final barcode diversity.

### Guide RNA design

In all experiments, VcCasTn gRNAs used 32 nt spacers and a 5’-CC Type IF PAM, while ShCasTn gRNAs used 23 nt spacers and a 5’-GTT Cas12k PAM. All gRNAs were designed to bind in the first half of the target CDS to ensure functional knockout by transposon insertion (Supplementary Table 3). Off-target potential was assessed using BLASTn (-dust no -word_size 4) of spacers against a local BLAST database created from all genomes present in an experiment, and spacers were discarded if off-target hits with E-value < 15 were identified. gRNAs with less seed region complementarity to off-targets were prioritized. Non-targeting gRNAs were designed by scrambling the spacer until no significant matches were found.

### Delivery methods

For natural transformation and electroporation, a culture of the community or isolate to be transformed was subcultured at OD_600_ = 0.2 and grown to OD_600_ = 0.5. In the case of the thiocyanate-degrading bioreactor in the absence of accurate OD measurements the culture was outgrown for two hours. For natural transformation 200 ng of vector harboring the *mariner* transposon (pHLL250^1^) for non-targeted insertion, or water for the negative control were added to 4 mL of OD_600_ = 0.5 outgrowth. Cultures were incubated for 3 hours shaking at 250 rpm at temperature appropriate for the isolate or community before being moved to the appropriate downstream analysis.

For electroporation, 20 mL of the community or isolate at OD_600_ = 0.5 was put on ice, centrifuged at 4,000*g* at 4°C for 10 minutes, and washed four times with 10 mL sterile ice-cold Milli-Q H_2_O. After a final centrifugation the pellet was resuspended in 100 μL of 2 ng/μL vector (pHLL250 or VcDART), or 100 μL of water as a negative control. This solution was then pipetted into a 0.2 cm gap ice-cold cuvette and electroporated at 3 kV, 200Ω, and 25 μF. The cells were immediately recovered into 10 mL of the community’s or isolate’s preferred medium and incubated shaking for 3 hours before being moved to the appropriate downstream analysis.

*E. coli* strain WM3064 containing the *mariner* transposon (pHLL250) for non-targeted editing, or the VcDART for targeted editing was cultured overnight in LB supplemented with carbenicillin (100 μg/mL) and DAP (60 μg/mL) at 37°C. Before conjugation the donor strain was washed twice in LB (centrifugation at 4,000*g* for 10 minutes) to remove antibiotics. Then, 1 OD_600_*mL of the donor was added to 1 OD_600_*mL of the recipient community or isolate and the mixture was plated on a 0.45 μm mixed cellulose ester membrane (Millipore) topping a plate of the recipient’s preferred media without DAP. In the case of the thiocyanate-degrading bioreactor, ~2 OD_600_*mL of the donor was added to 2 OD_600_*mL of the recipient community to ensure sufficient material despite the community’s slow growth. Plates were incubated at the ideal temperature for the recipient community or isolate for 12 hours before the growth was scraped off the filter into the media of the recipient community or isolate for downstream analysis.

### ET-Seq library preparation

The insertion junction sequencing library prep strategy for ET-Seq can be used (modification may be necessary) in any circumstance where high efficiency mapping of inserted DNA to a host loci is desired. For our purposes, DNA of the edited community or isolate was first extracted using the DNeasy PowerSoil Kit (QIAGEN). In the case of the nine-member community, 500 ng of DNA was used for both insertion junction sequencing and metagenomic library prep. For the SCN community, which had lower yields of DNA, 100 ng were used. As an internal standard, DNA from a previously constructed mutant library of *Bacteroides thetaiotaomicron* VPI-5482^5^, a species not present in the nine-member community or the thiocyanate-degrading bioreactor, was spiked into the community DNA at a ratio of 1/500 by mass. The *B. thetaiotaomicron* library had undergone antibiotic selection for its transposon insertions and was thus assumed to represent 100% transformation efficiency (i.e. every genome contained at least one mariner transposon insertion).

For metagenomic sequencing, library prep was conducted by the standard ≥100 ng protocol from the NEBNext Ultra II FS DNA Library Prep Kit for Illumina (NEB). For insertion junction sequencing, the same protocol was used with a number of modifications enumerated here (Extended Data Fig. 1). This insertion junction sequencing protocol has also been tested successfully with the ≤ 100 ng protocol of the NEBNext Ultra II FS DNA Library Prep Kit (NEB) and the KAPA HyperPlus Kit (Roche). For fragmentation an 8 minute incubation was used. A custom splinklerette adaptor was used during adaptor ligation to decrease non-specific amplification (Supplementary Table 4)^6,7^. For size selection 0.15X (by volume) SPRIselect (Beckman Coulter, Cat # B23318) or NEBNext Sample Purification Beads (NEB) were used for the first bead selection and 0.15X (by volume) were added for the second. From this selection, the DNA was eluted in 44 μL (instead of the suggested 15 μL) where it undergoes digestion before enrichment to cleave intact transposon delivery vector. All bead elutions were performed with Sigma Nuclease-Free water. pHLL250 underwent AseI digestion, while DART vectors underwent double digestion by AsiSI and SbfI-HF (NEB) (Supplementary Table 2). The DNA then underwent a sample purification using 1X AMPure XP beads (Beckman Coulter) to prepare it for PCR enrichment.

In PCR enrichment, the transposon junction was amplified by nested PCR. The PCRs followed the NEBNext Ultra II FS DNA Library Prep Kit for Illumina (NEB) PCR protocol, however in the first PCR the primers were custom to the transposon and the adaptor and the PCR was run for 25 cycles (Supplementary Table 4). The enrichment then underwent sample purification with a 0.7X size selection using SPRIselect or NEBNextSample Purification Beads from which 15 μL were eluted for the second PCR. This second PCR used custom unique dual indexing primers specific to nested regions of the insertion and adaptor and 6 cycles are used (Supplementary Table 4). Then another 0.7x size selection was conducted and the final library was eluted in 30 μL. Samples for metagenomic sequencing and insertion junction sequencing were then quality controlled and multiplexed using 1X HS dsDNA Qubit (Thermo Fisher) for total sample quantification, Bioanalyzer DNA 12000 chip (Agilent) for sizing, and qPCR (KAPA) for quantification of sequenceable fragments. Samples were sequenced on the iSeq100 or HiSeq4000 platforms.

### Genome sequencing, assembly, taxonomic classification, and database construction

For a full list of genome sequences used as read mapping references in this study see Supplementary Table 1. Assembly and annotation of genomes used as references for the SCN bioreactor experiment is described in Huddy et al.^8^. As the SCN bioreactor has been subjected to numerous genome-resolved metagenomic studies^8,9^ we endeavored to create a non-redundant database that contained all genomes previously observed in this reactor system. A set of 556 genomes assembled from this system were de-replicated at the species level using dRep v2.5.3^10^ with an average nucleotide identity (ANI) threshold of 95% and a minimum completeness of 60% as estimated by checkM v1.1.2^11^. A single genome representing each species level group was chosen by dRep based on optimizing genome size, fragmentation, estimated completeness, and estimated contamination resulting in 265 representative genomes. Genomes were taxonomically classified using GTDB-Tk^12^ with default options. Additionally, to display the taxonomic diversity of transformed organisms (Fig. 3b), a phylogenetic tree was constructed using ribosomal protein S3 (rpS3). Briefly, a custom Hidden Markov Model (HMM) was used to identify rpS3 sequences^13^ in the 265 representative genomes, and 262 rpS3 sequences were successfully identified. Sequences were aligned using muscle^14^, and a maximum likelihood phylogenetic tree was constructed using IQ-TREE^15^. Phylogenetic trees were pruned and annotated using iTol v5 (https://itol.embl.de/). To determine if an organism we identified as receiving exogenous DNA was ever previously isolated we used the ANI relative to the closest reference genome in the GTDB-Tk output. If one of our genomes had an ANI ≥ 95% relative to a known reference, and this reference genome was generated from an isolated bacterium, our target organism was considered to be previously isolated (Supplementary Table 1).

For 9-member community genomes assembled as part of this study, cultures were grown on R2A medium for 24 hours at 30°C and genomic DNA was extracted with the DNeasy Blood and Tissue DNA Kit (Qiagen) with pre-treatment for Gram-positive bacteria. Genomic DNA was sheared mechanically with the Covaris S220 and processed with the NEBNext DNA Library Prep Master Mix Set for Illumina (NEB) before submitting for sequencing on an Illumina MiSeq platform generating paired end 150 bp reads. Raw sequencing reads were processed to remove Illumina adapter and phiX sequence using BBduk with default parameters, and quality trimmed at 3’ ends with Sickle using default parameters (https://github.com/najoshi/sickle). Assemblies were conducted using IDBA-UD v1.1.1^16^ with the following parameters: –pre_correction –mink 30 – maxk 140 –step 10. Following assembly, contigs smaller than 1 kbp were removed and open reading frames (ORFs) were then predicted on all contigs using Prodigal v2.6.3^17^. 16S ribosomal rRNA genes were predicted using the 16SfromHMM.py script from the ctbBio python package using default parameters (https://github.com/christophertbrown/bioscripts). Transfer RNAs were predicted using tRNAscan-SE^18^. The full metagenome samples and their annotations were then uploaded into our in-house analysis platform, ggKbase, where genomes were manually curated via the removal of contaminating contigs based on aberrant phylogenetic signatures (https://ggkbase.berkeley.edu).

For each ET-Seq experiment a genomic database is constructed using the ETdb component of the ETsuite software package. Each database contains the nucleotide sequences of the expected organisms in a sample, any vectors used, any conjugal donor, and the spike in control organism. Briefly, all genomic sequences are formatted into a bowtie2 index to allow read mapping, a tabular correspondence table between all scaffold names and their associated genome is constructed (scaff2bin.txt), and a table (genome_info.txt) of standard genomic statistics is calculated including genome size, GC content, and number of scaffolds. Following database construction, a label is manually added to each entry in the genome info table to indicate if the entry represents a target organism, a vector, or a spike in control organism. All data are propagated into a single folder that can be used by the ETmapper software for downstream mapping and analysis.

### Identification and quantification of insertion junctions and barcodes

To identify and map transposon insertion junctions and their associated barcodes in a mixed population of microbial cells, reads (150 bp X 2) generated from PCR amplicons of putative transposon insertion junctions are first processed using the ETmapper component of the ETsuite software package implemented in R with the following steps: First reads are quality trimmed at the 3’ end to remove low quality bases (Phred score ≤ 20) and sequencing adapters using Cutadapt v2.10^19^. Cutadapt is then used to identify and remove provided transposon model sequences from the 5’ end of forward reads, requiring a match to 95% of the shortest transposon sequence in a provided set and allowing a 2% error rate. Read pairs where no transposon model sequence is identified in the forward read are discarded. All identified and trimmed transposon models are paired with their respective reads, stored, and barcodes are identified in these sequences by searching for a known primer binding site sequence flanking the 5’ end of the barcode (5’-CTATAGGGGATAGATGTCCACGAGGTCTCT-3’) allowing for 1 mismatch. Subsequently, the 20 bp region following the known primer binding site is extracted as the barcode sequence and associated with its respective read. The 3’ end of the paired reverse reads are then trimmed to remove any transposon model sequence using Cutadapt, and only read pairs where one mate is at least ≥ 40 bp following all trimming are retained for downstream mapping and analysis. The fully trimmed paired end reads now consisting of only genomic sequence following the transposon insertion site are mapped to the ETdb database used in a given experiment using bowtie2 with default options^20^. Mapped read files are converted into a hit table indicating the mapped genome, scaffold, genomic coordinates, mapQ score, and number of alignment mismatches for each read in a pair using a custom Python script, bam_pe_stats.py, provided with ETsuite. This table is then merged with read-barcode assignments to generate a final hit table with the mapping information about each read pair, the transposon model identified, and the associated barcode found for that read pair. Finally mapped read pairs are only retained for downstream quantification if both reads map to the same genome, at least one mapped read in a pair has a mapQ score ≥ 20, and a barcode was successfully identified and associated with the read pair.

To quantify the number of unique barcodes and their associated reads mapping to organisms in each sample of an experimental run, the filtered hit tables were processed using the ETstats component of the ET-Seq software package with the following steps: Initially, all barcodes identified across all samples in an experiment are aggregated and clustered using Bartender^21^ with the following supplied options: −l 4 −s 1 −d 3. Barcode clusters and their associated barcodes/reads were only retained if all of the following criteria were true: (1) ≥ 75% of the reads in a cluster mapped to one genome (the majority genome), (2) ≥ 75% of the reads in a cluster were associated with the same transposon model (the majority model), and (3) the barcode cluster had at least 2 reads. Subsequently, when quantifying reads and barcodes in each sample of an experiment, the genome a read was mapped to and the transposon model it was associated with had to agree with the majority assignments for the barcode cluster assigned to that read’s barcode to be counted. Finally, we were aware that Illumina patterned flow cell related index swapping would result in reads from a barcode cluster being misassigned across samples, even when using unique dual indexing^22^. We could not simply limit barcode clusters to be associated with only one sample, as our spike in control organisms contain the same pool of barcodes and are added to every sample. Thus we estimated an empirical index swap rate across each experiment and required that the number of reads (X) for a barcode to be positively identified in a sample be always ≥ 2 and ≥ the binomial mean of observed read counts expected in any sample for a barcode cluster with (R) reads across (N) samples based on the estimated swap rate (S) + 2 standard deviations (**Eqn. 1**)

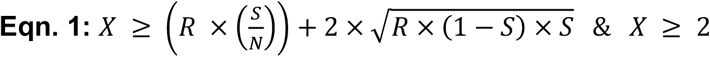

The index swap rate for an experiment was empirically estimated from barcode clusters assigned only to target organisms based on the assumption that it would be highly unlikely for a barcode cluster to have truly originated from independent integration events into the same organism in more than one sample. Thus we assumed that for each barcode cluster associated with target organisms, the majority of reads originated from the true sample and reads assigned to other samples represented swaps. This is opposed to barcode clusters associated with our spike-in organism, conjugal donor organism, or vectors which contain the same pool of barcodes directly added to multiple samples. To identify swapped read counts we first quantify the total count of all reads assigned to the majority genome across barcode clusters but that are not associated with the majority sample of that cluster (E). Then we quantify the total count of reads associated with the majority genome and associated with the majority sample across all clusters (C). Then experiment wide swap rate was estimated by dividing the total number of reads not associated with majority samples by the total number of reads (**Eqn. 2**)

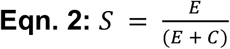

Following filtering, a hit table is returned that indicates for each genome in each sample, the number of unique barcode clusters that were recovered, and the total number of reads associated with these barcodes. As a final check for false positives during ET-Seq development we included an organism genome as an ETsuite mapping target, *Sinorhizobium meliloti*, which was not physically included in our 9-member synthetic or thiocyanate-degrading communities. We did not detect any barcodes or reads associated with this genome.

### Metagenomic data processing and coverage calculation

Each ET-Seq sample is split and in parallel undergoes shotgun metagenomic sequencing to determine the relative quantities of organisms present in the sample at the time of sampling. Raw read files from metagenomic data are also processed using the ETmapper component of the ETsuite software package with the following steps: First reads are quality trimmed at the 3’ end to remove low quality bases (Phred score ≤ 20) and sequencing adapters using Cutadapt v2.10^19^. Read pairs where at least one mate is not ≥ 40 bp in length are discarded. Trimmed read pairs are mapped to the ETdb database used in a given experiment using bowtie2^20^ with default parameters. Mappings are filtered to require a minimum identity ≥ 95% and minimum mapQ score ≥ 20, and coverage is calculated using a custom script, calc_cov.py, included with the ETsuite software.

Metagenomic sequencing for one biological replicate of the thiocyanate-degrading bioreactor community (Conjugation - Control Sample - Replicate 3) failed. Metagenomic coverage values for this replicate were generated by averaging the values from the other two biological replicates.

### ET-Seq normalization and calculation of insertion efficiency

To account for differences in sequencing depth, transposon junction PCR template amount, and relative abundance of microbes in a community the data generated from both ET-Seq and shotgun metagenomics were each normalized independently to values from the spike in control organism, *B. thetaiotaomicron*, and then ET-Seq data is subsequently normalized by metagenomic abundance as follows: Initially read count tables from ET-Seq and metagenomics are filtered to remove any ET-Seq read count associated with < 2 barcodes and any metagenomic read count < 10 reads. Next a size factor for each sample is calculated based on the geometric mean of *B. thetaiotaomicron* reads for ET-Seq samples and *B. thetaiotaomicron* coverage for metagenomics samples. ET-Seq read counts and metagenomic coverage values are then divided by their respective sample size factors to create normalized values. Normalized ET-Seq read counts are then divided by their paired normalized metagenomic coverage values to generate ET-Seq read counts that are fully normalized to both ET-Seq sequencing depth and metagenomic coverage. Finally fully normalized ET-Seq read counts for target organisms are divided by the fully normalized ET-Seq read count of *B. thetaiotaomicron* from an experiment (a constant that represents the number of reads that would be obtained from an organism with 100% of its chromosomes carrying insertions). The resulting values for each target organism in a sample represent an estimate of the fraction of that organism’s population that received insertions (Insertion Efficiency). Additionally, we multiply a target organism’s insertion efficiency by the fractional relative abundance of that organism in a sample, based on metagenomic data, to estimate the fraction of an entire sample population that is made up of cells of a given species that received insertions (Insertion-Receiving Fraction in Total Community).

### ET-Seq validation and establishing limits of detection and quantification

To validate ET-Seq and establish both a limit of detection (LOD) and limit of quantification (LOQ) for the assay, a library of *K. michiganensis* transposon mutants was constructed by antibiotic selection following conjugation with pHLL250 (as described above), and this library was added to untransformed samples of the combined nine-member community to create a transformed cell concentration gradient. Technical triplicate samples were created where 1%, 0.1%, 0.01%, 0.001% and 0% of the total *K. michiganensis* cells (by OD_600_) in the mixture were those derived from the transformed library. All samples (n = 15) were subjected to ET-Seq (as described above), and pooled samples across all concentrations for each technical triplicate (n = 3; 5 concentrations) were analyzed for community composition using shotgun metagenomics (as described above). ET-Seq insertion efficiencies and insertion-receiving fraction in total community values were averaged across technical replicates. Additionally, to derive the fraction of transformed *K. michiganensis* cells that made up the total community (not just the *K. michiganensis* sub-population), the known fraction of *K. michiganensis* cells that were transformed in a sample was multiplied by the measured relative abundance of *K. michiganensis* in a given technical replicate, and these values were averaged across technical replicates.

To derive the LOD and LOQ for ET-Seq a linear regression was performed using the lm function in the base package of R^23^ using the known fraction of transformed *K. michiganensis* cells that made up the total community as the independent variable and the ET-Seq estimated per community insertion efficiency as the dependent variable. The sample where transformed *K. michiganensis* made up 0% of the community was not included in the regression analysis, but was reserved to demonstrate zero response with no transformed cells present. LOD was calculated as 3.3 * standard error of the regression / slope. The LOQ was calculated as 10 * standard error of the regression / slope.

### Identification of positive transformations and statistical analysis

For all ET-Seq experiments conducted we initially determined if any ET-Seq estimated per community insertion efficiency was larger than the LOD. Values larger than the LOD constituted a positive detection. For comparative statistical analysis conducted to compare insertion efficiencies between transformation methods (Fig. 2b) only values that had a corresponding insertion-receiving fraction in total community > LOQ were used. Statistical testing was conducted using Analysis of Variance (ANOVA) implemented in the aov function in R^23^. Post-hoc testing was conducted using the TukeyHSD function in R. Traditional 95% confidence intervals were calculated using the groupwiseMean function of the rcompanion package in R.

### Multiple delivery experiments in communities

To test multiple delivery methods on the nine-member community, all members were grown at 30°C with *Bacillus sp. AnTP16* and *Methylobacterium sp. UNC378MF* in R2A liquid media while all other members were inoculated in LB. Equal amounts of community members were then combined by OD_600_. This consortium then underwent transformation (of pHLL250), conjugation (pHLL250 in WM3064), and electroporation of the pHLL250 vector (described in Delivery Methods section). After delivery the community was spun down at 5,000*g* for 10 minutes, washed once with LB and then spun down and frozen at −80°C until genomic DNA extraction.

The thiocyanate-degrading microbial community was sampled for delivery testing from biofilm on a four liter continuously stirred tank reactor that had been maintained at steady state for over a year. The reactor is operated with a two day hydraulic residence time, sparged with laboratory air at 0.9 L/min, and fed with a mixture of molasses (0.15% w/v), thiocyanate (250 ppm), and KOH to maintain pH 7. OD measurements were not feasible on the biofilm so we used its wet mass to approximate equivalent OD and thus cell numbers to those used for the nine-member community. This community underwent the same transformation, electroporation, and conjugation delivery approaches as the nine-member community, however in all steps requiring media, LB was replaced with molasses media (no thiocyanate). After delivery the community was spun down at 5,000*g* for 10 minutes, washed once with molasses media and then spun down and frozen at - 80°C until genomic DNA extraction.

### Benchmarking DART systems in *E. coli*

We first constructed several DART systems to identify variants capable of efficient transposition by conjugative delivery to *E. coli*. We performed parallel conjugation of each DART vector variant containing Gm^R^ Tn cargo (2.1 kbp) and either a non-targeting gRNA or one of two *lacZ*-targeting gRNAs for each system. For VcDART, variation of the promoter controlling the expression of VcCasTn components did not significantly impact transposition efficiency (Extended Data Fig. 4c-d). Similarly for ShDART, expression of the sgRNA in three distinct transcriptional configurations did not significantly impact transposition efficiency (Extended Data Fig. 4e-f). Since promoter and transcriptional configuration variation had insignificant effects on transposition efficiency--and to remove the requirement for promoter induction and reliance on T7 RNA polymerase--we performed target specificity benchmarking of VcDART and ShDART using the same constitutive P_lac_ promoter. In this experiment, ShDART Cas and Tns genes and sgRNA were encoded in the original transcriptional configuration and under control of the same promoter in which ShCasTn was first characterized by Strecker et al.^3^.

The *lacZ*-targeting gRNAs were designed to target the *lacZ* α-peptide present in the conjugation recipient strain *E. coli* BL21(DE3) but absent in the *lacZ*ΔM15 strains used as cloning host (*E. coli* EC100D-*pir*+) or conjugation donor (*E. coli* WM3064), preventing transposition until delivery into the recipient cell (Extended Data Fig. 4a). Donor WM3064 strains were transformed and cultivated as described above, and recipient BL21(DE3) was inoculated from glycerol stock into 100 mL LB in a 250 mL baffled shake flask at 37°C 250 rpm. Conjugations were performed as described above using LB medium and 37°C incubation for every step, except that 0.1 mM IPTG was added to VcDART conjugation plates in Extended Data Fig. 4d to induce transcription from P_T7-lac_ and T7 RNA polymerase expression in *E. coli* BL21(DE3). Transposition efficiencies were calculated as the percentage of colonies resistant to 10 μg mL^-1^ gentamycin relative to viable colonies in absence of gentamycin.

On/off-target analysis was performed for one *lacZ*-targeting guide for each DART system by outgrowth under selection followed by genomic DNA extraction and ET-Seq. Specifically, approximately 10,000 transconjugant cfu were plated on LB agar with gentamycin, incubated at 37°C overnight, scraped from agar into liquid LB medium, diluted to OD_600_ = 0.25 into 10 mL LB plus gentamycin in 50 mL conical tubes, incubated at 37°C 250 rpm until OD_600_ = 1.0, centrifuged at 4,000*g*, and frozen for downstream analysis. To determine the percent of selectable transposed colonies possessing on-target and off-target edits, the total number of selectable colonies was adjusted (Extended Data Fig. 4b) for on-target and off-target percent as determined by ET-Seq (Fig. 4b). ET-Seq analysis was conducted on triplicate platings of DART transconjugants (n = 3 for each system) to identify transposon insertion locations and quantify on-target vs. off-target insertions. As the targeted genomic region encoding the *lacZ* α-peptide is duplicated in *E. coli* BL21 (DE3), one of the two duplicated regions (749,903 bp --> 750,380 bp) was removed prior to analysis to allow unambiguous mapping assignment. Subsequently, the standard ETsuite analysis pipeline (as described above) was used to identify and map 300 bp X 2 reads containing transposon junctions back to the recipient BL21(DE3) genome and cluster barcodes that corresponded to unique insertion events. To confirm an insert location we first identified the exact transposon-genome junction mapping coordinate that was the most frequent in the reads of a barcode cluster (prime location) then required that a barcode cluster had: (1) at least 75% of its reads coming from within 3 bp of the prime location and (2) at least 75% of its reads mapping to the same strand. If these criteria were true the barcode cluster was counted as a unique insertion and the prime location was used as the mapping locus by ET-Seq. An on-target insertion was evaluated as a barcode cluster with a prime location within 200 bp downstream of the 3’ end of the protospacer target. Finally all distances reported from the protospacer target site were calculated from the last base pair of the 3’ end of the protospacer.

### VcDART-mediated targeted editing in a community

VcDART vectors encoding constitutive VcCasTn, constitutive *bla:aadA* Tn cargo (2.7 kbp), and either a non-targeting (pBFC0888), *K. michiganensis* M5aI *pyrF*-targeting (pBFC0825), or *P. simiae* WCS417 *pyrF*-targeting (pBFC0837) constitutive crRNA were transformed into *E. coli* WM3064. Conjugations of these vectors into the nine-member community were performed as described above on filter-topped LB agar plates with 12 hr incubation at 30°C. Lawns were scraped from filters into 10 mL LB medium, vortexed, and 1 OD_600_*mL from each lawn was plated on LB agar supplemented with 1 mg mL^-1^ 5-FOA, 100 μg mL^-1^ carbenicillin, 100 μg mL^-1^ streptomycin, and 100 μg mL^-1^ spectinomycin. Following 3 days of incubation at 30°C, all cells were scraped from the agar into 10 mL R2A medium, vortexed, diluted into 10 mL R2A supplemented with 20 mg mL^-1^ uracil (for no selection controls) or R2A with uracil, 5-FOA, carbenicillin, streptomycin, and spectinomycin to OD_600_ = 0.02, and split evenly across 4 wells (2.5 mL/well) of a 24 deep well plate. After cultivation at 30°C and 750 rpm for 1 week, only the cultures conjugated with VcDART containing *pyrF*-targeting crRNA had grown in presence of antibiotics and 5-FOA. A small portion of each of these cultures was serially diluted in R2A and plated on LB agar plus antibiotics to isolate and assay colonies by targeted PCR and Sanger sequencing of *pyrF* loci. The remainder of each culture was centrifuged at 4,000*g* for 10 min and frozen at −80°C for downstream bacterial 16S rRNA V4 amplicon metagenomic sequencing (Novogene). Relative abundances were calculated as described below (16S rRNA V4 amplicon analysis) for pre-conjugation nine-member community cultures and post-selection *pyrF*-targeted cultures.

### 16S rRNA V4 amplicon analysis

16S rRNA V4 amplicon sequencing was conducted using the 515F (5’-GTGCCAGCMGCCGCGGTAA-3’) and 806R (5’-GGACTACHVGGGTWTCTAAT-3’) universal bacterial primer set to generate 250 bp x 2 reads (Novogene). Samples were processed using the UPARSE pipeline within the USEARCH software package to merge read pairs, remove primers, quality filter sequences, remove chimeras, identify unique sequence variants (ZOTUs), and quantify their abundance across samples as described previously^24^. To assign ZOTU sequences to species known to be in our community mixture, we queried all 890 identified ZOTUs against a custom database of 16S sequences derived from the genomes of the nine-member community constituents using USEARCH^25^ ZOTUs with 100% identity to a 16S sequence in our database were assigned to the matched species, and all matches < 100% identity were counted as “Other”. Counts coming from ZOTUs of the same taxonomic assignment were merged and the relative abundance of a species was calculated as its read counts divided by the total read counts for the sample.

### Statistics and reproducibility

All transformations (natural transformation, conjugation, electroporation) and subsequent analyses were performed for three independent replicates.

### Reporting summary

Further information on research design is available in the Nature Research Reporting Summary linked to this paper.

### Data availability

Summary data for genomes, plasmids, and oligonucleotides used in this study can be found in Supplementary tables 1-4. Sequence data for all genomes assembled as part of this study and newly constructed plasmids are in submission to NCBI with accession numbers pending. Sequence data for genomes taken from Huddy, et al^8^ are in submission to NCBI with accession numbers pending. Sequence data for genomes taken from Kantor, et al^9^ are available under NCBI BioProject accession no. PRJNA279279. All genomes and plasmids used in the project will also be made available on ggKbase (https://ggkbase.berkeley.edu/). Raw count data for all experiments including both metagenome and ET-seq information is available at https://github.com/SDmetagenomics/ETsuite/tree/master/manuscript_data.

### Code availability

Custom R scripts for ET-Seq analysis and code used in the construction of figures are available at https://github.com/SDmetagenomics/ETsuite.

## Acknowledgments

We thank Morgan N. Price for data analysis input, Patrick Pausch for experimental advice, Shana L. McDevitt, Eileen Wagner, and Hitomi Asahara for help with sequencing, and Trent R. Northen for directional advice. Funding was provided by m-CAFEs Microbial Community Analysis & Functional Evaluation in Soils, a project led by Lawrence Berkeley National Laboratory supported by the U.S. Department of Energy, Office of Science, Office of Biological & Environmental Research under contract number DE-AC02-05CH11231. Support was also provided by the Innovative Genomics Institute at UC Berkeley. B.E.R. and B.F.C. are supported by the National Institute of General Medical Sciences of the National Institute of Health under award numbers F32GM134694 and F32GM131654. Schematics were created with BioRender.com.

## Contributions

B.E.R., S.D., B.F.C, A.M.D., J.F.B, and J.A.D. conceived the work and designed the experiments. B.E.R., B.F.C., C.H., M.X., Z.Z., D.C.S., K.T., T.K.O., and N.K. conducted the molecular biology included. S.D., A.C.-C., C.H., and R.S. developed the bioinformatic analysis. B.E.R., S.D., B.F.C., A.M.D., J.F.B., and J.A.D. analyzed and interpreted the data.

## Additional Information

## Extended Data

**Extended Data Fig. 1.**
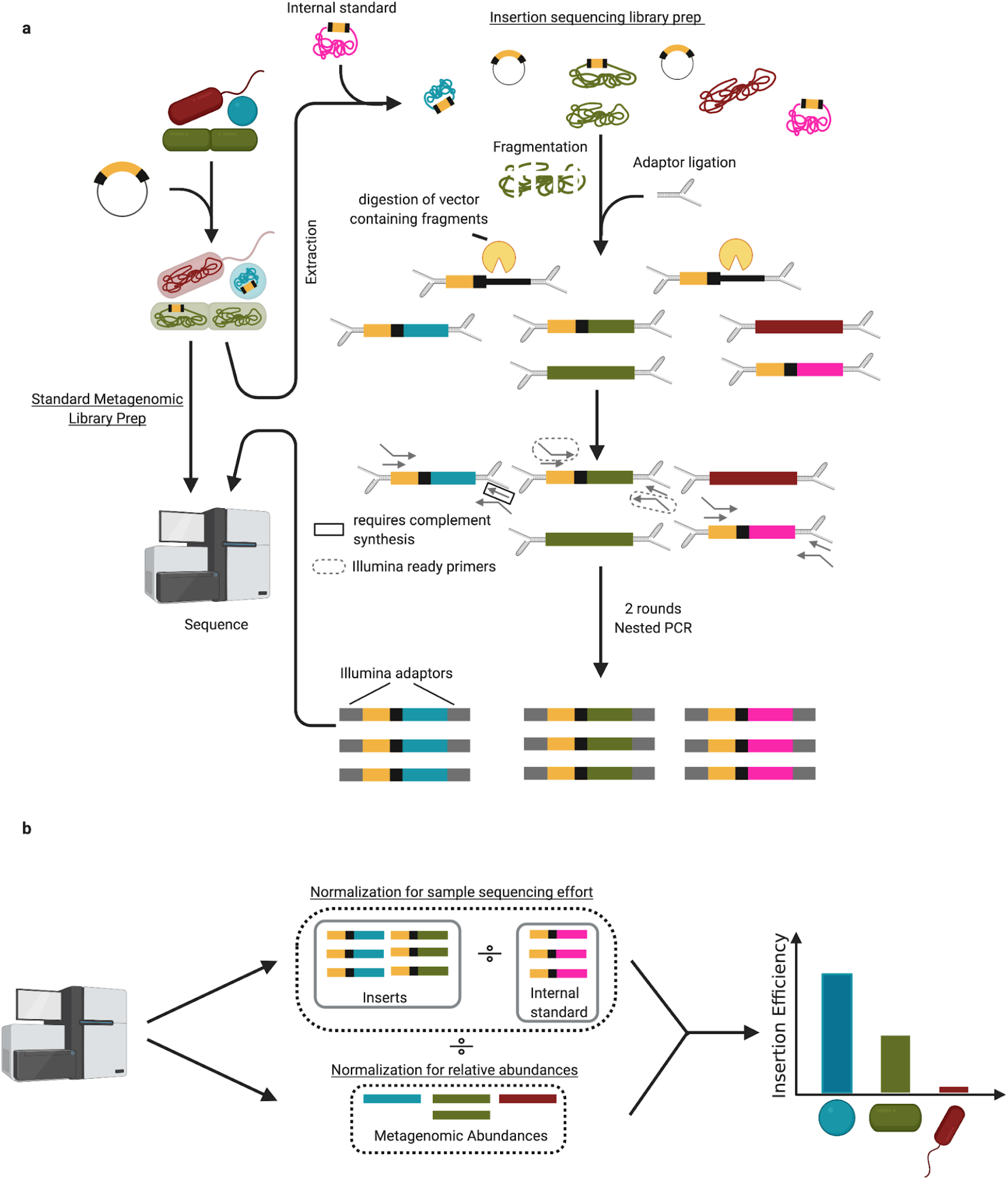
Library preparation and data normalization for ET-Seq. **a**, ET-Seq requires low-coverage metagenomic sequencing and customized insertion sequencing. Insertion sequencing relies on custom splinkerette adaptors, which minimize non-specific amplification, a digestion step for degradation of delivery vector containing fragments, and nested PCR to enrich for fragments containing insertions with high specificity. The second round of nested PCR adds unique dual index adaptors for Illumina sequencing. **b**, This insertion sequencing data is first normalized by the reads to internal standard DNA which is added equally to all samples and serves to correct for variation in reads produced per sample. Secondly, it is normalized by the relative metagenomic abundances of the community members.

**Extended Data Fig. 2.**
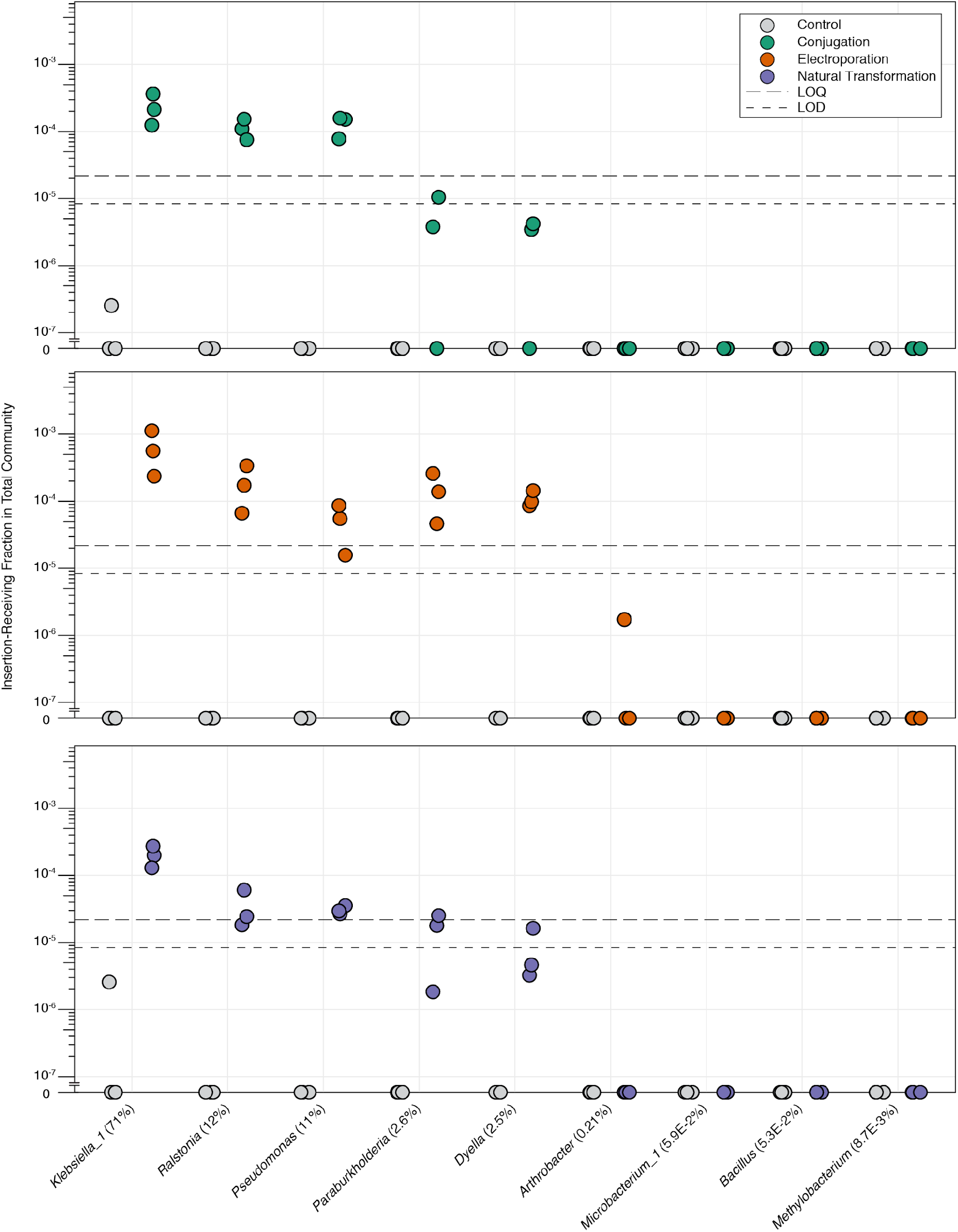
ET-Seq determined insertion efficiencies for all nine consortium members as a fraction of the entire community. ET-Seq determined insertion efficiencies for conjugation, electroporation, and natural transformation on the nine-member synthetic community (n = 3 biological replicates). The values shown are the estimated fraction a constituent species’s transformed cells make of the total community population. Control samples received no exogenous DNA. Average relative abundance across all samples is indicated in parentheses (n = 18 independent samples). LOD and LOQ are indicated in plots by short and long dashed lines respectively.

**Extended Data Fig. 3.**
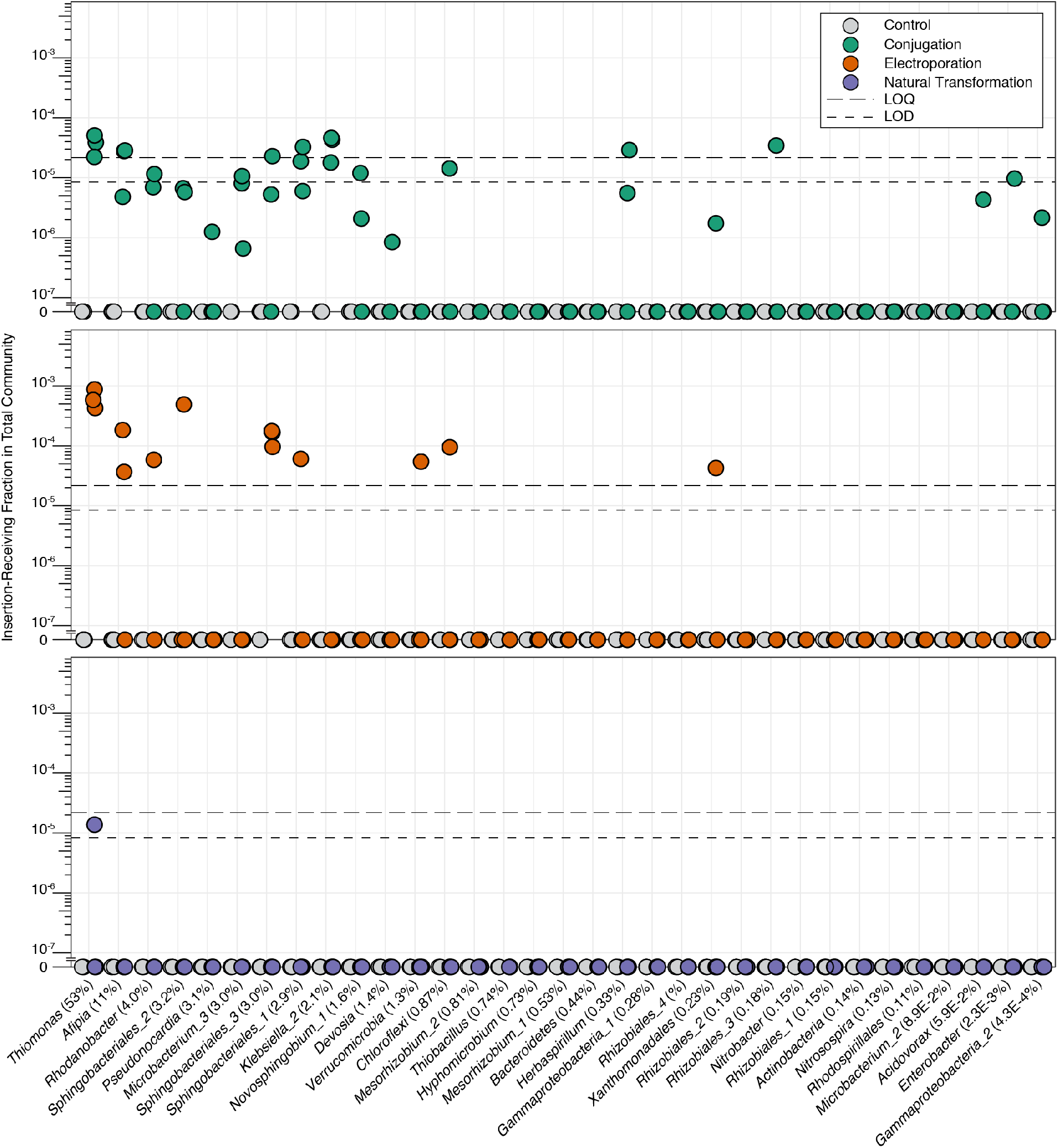
ET-Seq determined insertion efficiencies for all thiocyanate-degrading bioreactor community members as a fraction of the entire community. ET-Seq determined insertion efficiencies for conjugation, electroporation, and natural transformation on the thiocyanate-degrading bioreactor community (n = 3 biological replicates). The values shown are the estimated fraction a constituent species’s transformed cells make of the total community population. Control samples received no exogenous DNA. Average relative abundance across all samples is indicated in parentheses (n = 17 independent samples; due to a single failed metagenomic sequencing replicate, see methods). LOD and LOQ are indicated in plots by short and long dashed lines respectively.

**Extended Data Fig. 4.**
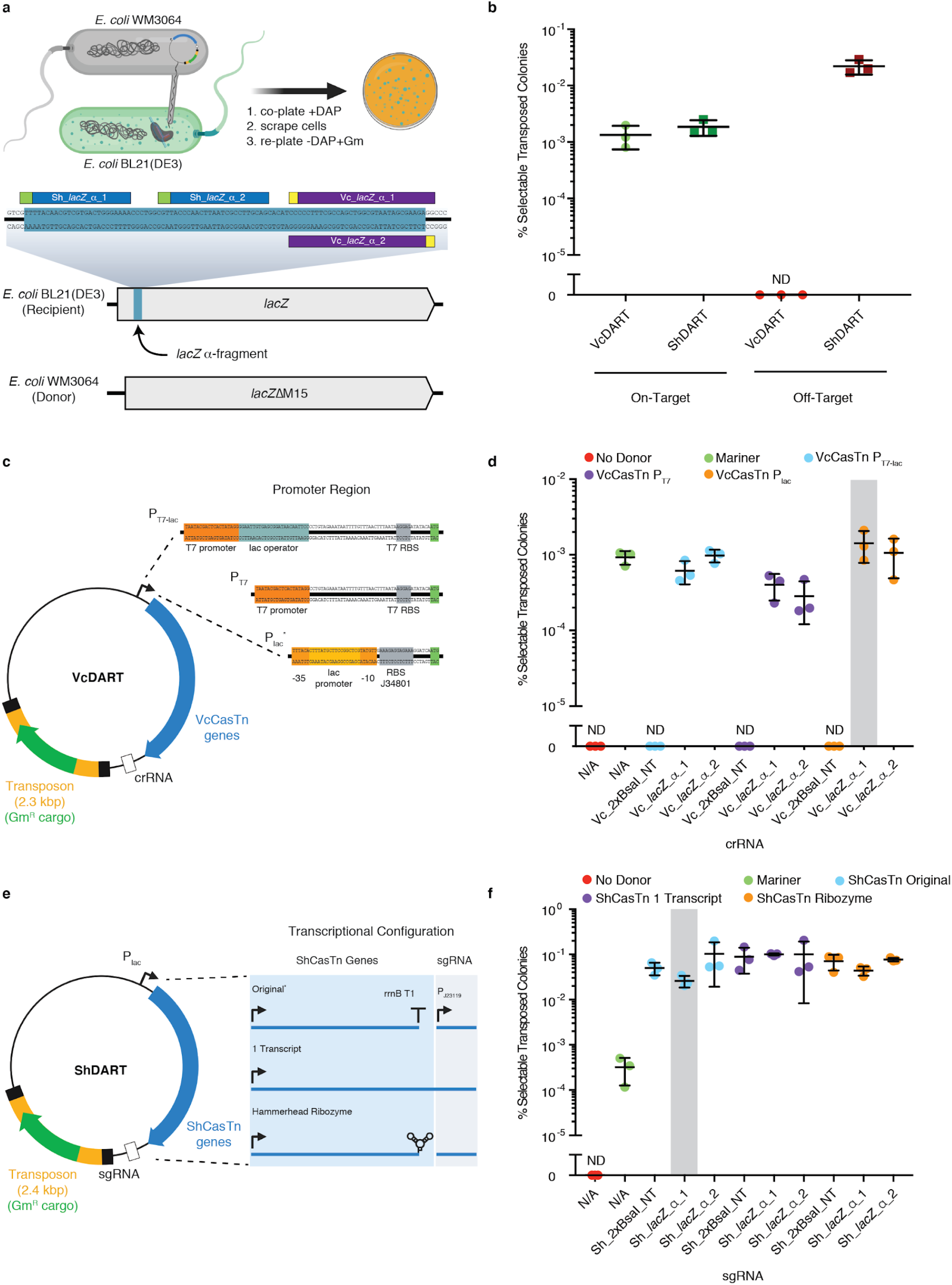
Benchmarking DART vectors. **a,** *E. coli* WM3064 to *E. coli* BL21(DE3) conjugation, transposition, and selection schematic (top) and guide RNAs targeting the *lacZ* α-fragment of recipient BL21(DE3), which is absent from donor WM3064 (bottom). **b,d,f,** Percent selectable transposed colonies is calculated as the number of colonies obtained with gentamycin selection divided by total viable colonies in absence of selection. **b,** Insertion receiving colonies divided into on- and off-targeted. This was calculated by multiplying % selectable colonies for representative guides in **d** and **f** (highlighted by grey bars) by the on- or off-target rates (shown in Fig. 4). **c**, Transposition with VcDART was tested with three promoters. The variant using the P_lac_ promoter, harvested from pHelper_ShCAST_sgRNA^18^, was also used for Fig. 4, 5, and Extended Data Fig. 4b (*). **d,** Efficiencies of VcDART using various promoters. **e,** Transposition with ShDART was tested with three transcriptional configurations, all using P_lac_^18^. The configuration used for characterization of ShCasTn originally^18^ was also used for Fig. 4 and Extended Data Fig. 4b (*). **f,** Efficiencies of ShDART using various promoters. **b, d, f,** Crossbar indicates mean and error bars indicate one standard deviation from the mean (n = 3 biological replicates). Guide RNAs ending in “NT” are non-targeting negative control samples.

